# The Relationship between BOLD and Neural Activity Arises from Temporally Sparse Events

**DOI:** 10.1101/644419

**Authors:** Xiaodi Zhang, Wen-Ju Pan, Shella Dawn Keilholz

**Author notes:** Corresponding author. (Shella Keilholz). E-mail addresses (Xiaodi Zhang), (Wen-Ju Pan).

## Abstract

Resting state functional magnetic resonance (rs-fMRI) imaging offers insights into how different brain regions are connected into functional networks. It was recently shown that networks that are almost identical to the ones created from conventional correlation analysis can be obtained from a subset of high-amplitude data, suggesting that the functional networks may be driven by instantaneous co-activations of multiple brain regions rather than ongoing oscillatory processes. The rs-fMRI studies, however, rely on the blood oxygen level dependent (BOLD) signal, which is only indirectly sensitive to neural activity through neurovascular coupling. To provide more direct evidence that the neuronal co-activation events produce the time-varying network patterns seen in rs-fMRI studies, we examined the simultaneous rs-fMRI and local field potential (LFP) recordings in rats performed in our lab over the past several years. We developed complementary analysis methods that focus on either the temporal or spatial domain, and found evidence that the interaction between LFP and BOLD may be driven by instantaneous co-activation events as well. BOLD maps triggered on high-amplitude LFP events resemble co-activation patterns created from rs-fMRI data alone, though the co-activation time points are defined differently in the two cases. Moreover, only LFP events that fall into the highest or lowest thirds of the amplitude distribution result in a BOLD signal that can be distinguished from noise. These findings provide evidence of an electrophysiological basis for the time-varying co-activation patterns observed in previous studies.

## 1. Introduction

Functional magnetic resonance imaging (fMRI) is a noninvasive method that uses the blood oxygenation level-dependent (BOLD) (1) signal to measure the neural activity in different parts of the brain. In resting state fMRI (rs-fMRI) (2), a statistical relationship between the spontaneous activity of different areas of the brain in the absence of an explicit stimulation or task indicates that the regions are functionally related (2, 3). Temporal correlation is often used to measure this functional connectivity, and the spatial patterns of the BOLD signal correlation show high similarity to many established brain networks, including motor, visual, language, default mode, and attention networks (2, 4-8). However, as an imaging method, rs-fMRI inevitably suffers from physical limitations on its temporal resolution (∼1s), as well as the fact that neural activity is measured indirectly though neurovascular coupling (9).

Recent rs-fMRI studies show that the BOLD information is compressed into the relatively few discrete events that have the highest amplitudes. Both co-activation patterns (10) and point process methods (11) can extract information from those few selected time frames. The results closely resemble the spatial patterns of the cross correlation maps obtained from the entire dataset, which suggests that the BOLD interregional correlation is driven by the discrete, high amplitude events.

While these approaches that focus on discrete BOLD events instead of a continuous interaction between brain regions have provided new insights into brain function, they rely on an indirect measurement of neural activity. Extracellular local field potential (LFP) measurements, on the other hand, are not only a direct measure of neural activity, but also provide high temporal resolution (∼1ms), albeit from a limited number of sites (12). In this paper, we investigate two key questions about the neural basis of rs-fMRI: 1) Do the BOLD events that are the basis of co-activation patterns reflect distinct, high amplitude neural events? 2) Is there a range of neural activity which reliably produces a BOLD response that can be distinguished from noise?

Most studies that analyze the LFP-BOLD relationship use temporal correlation (9, 12-14) or mutual information (14), which are not well suited to the analysis of distinct events. Inspired by the idea of the spike-triggered average (15) and co-activation patterns (CAPs) (10), we propose two new methods called the BOLD-triggered average of LFP power (referred to as the BOLD-triggered average in the rest of the article) and LFP-triggered co-activation patterns (LFP-CAPs) to see if the relationship between LFP and BOLD is driven by discrete high-amplitude events. The BOLD-triggered average shows the average time course of LFP power preceding high amplitude BOLD events, whereas the LFP-CAPs average the fMRI frames a certain lag (depending on the anesthetic agents) after high amplitude broadband LFP events to show the spatial distribution of the brain regions that “co-activate” with the neuronal activity recorded by LFP. The two methods provide complementary information in both the temporal and spatial domains, and the results suggest that the relationship between LFP and BOLD is also driven by a few distinct events that have the highest amplitudes in either the BOLD or LFP power. Further analysis shows that those high LFP events can be classified into several groups, producing spatial patterns that are similar to those obtained from the original CAPs method, which solely uses rs-fMRI data.

## 2. Results

### 2.1. BOLD-triggered Average of LFP Power Time Courses

The spike-triggered average is widely used for the analysis of electrophysiological data. Each action potential produced by the neuron is considered as an event, which triggers the extractions of the stimulus in a short time window preceding the action potential event. Though the stimuli in the individual windows appear random, the averaged stimulus across all action potential events typically exhibits a pattern that is likely to cause the firing of the neuron, and thus correspond to the receptive field of the neuron. We hypothesize that, if we separate the BOLD time points (BOLD events) into several groups based on their amplitudes, and within each group, average the LFP power time course preceding the BOLD events, we may get a pattern in LFP power (both in temporal domain and frequency domain) that is likely to cause the occurrence of a BOLD event with a specified amplitude.

To test our hypothesis, we analyzed 337 simultaneous single slice fMRI and primary somatosensory cortex (S1) LFP recordings from both hemispheres from 36 Sprague–Dawley rats (male, 200–300 g, Charles River) under isoflurane (ISO. n = 100) ranging from 1% to 2%, or dexmedetomidine (DMED, n = 237) anesthesia on a 9.4T/20 cm horizontal bore small animal MRI system (Bruker, Billerica, MA). The details of animal preparation and parameter settings for data acquisition are described *in section 5. Materials and Methods*. A subset of the data was selected for further analysis if it met the following criteria: having high LFP broadband power versus BOLD S1-seed cross-correlation, and a bilateral symmetry in BOLD S1-seeded correlation map (to ensure a normal inter-hemisphere functional connectivity). By these criteria, 22 scans under DMED and 32 scans under various ISO concentrations were selected out of 337 scans. The data and code will be available upon request.

The BOLD signal was extracted from the ROI that has the highest correlation between LFP power and BOLD, which is found near the tip of the electrode. Then both the BOLD signal and LFP broadband power were z-scored and pooled together for by anesthesia (ISO and DMED). Within each dataset, the BOLD signal time points were evenly divided into 10 percentile groups based on their amplitudes. Figure 1 illustrates the process of obtaining the BOLD-triggered average time course. First the percentiles were calculated from the pooled distribution to obtain the thresholds for each percentile group (shown in the color-coded histogram), and the time points within the thresholds were selected as the triggers. Then the LFP broadband power time course preceding each BOLD trigger was extracted and averaged across all fMRI scans, and the resulting time course is referred to as the “BOLD-triggered average”. The adjacent triggers are considered as separated triggers, though alternatively one could make them become a single trigger, weighted by the duration of the event. The two methods produce very similar results (see Appendix, Figure A.1); we used the former for the rest of the paper.

**Figure 1.**
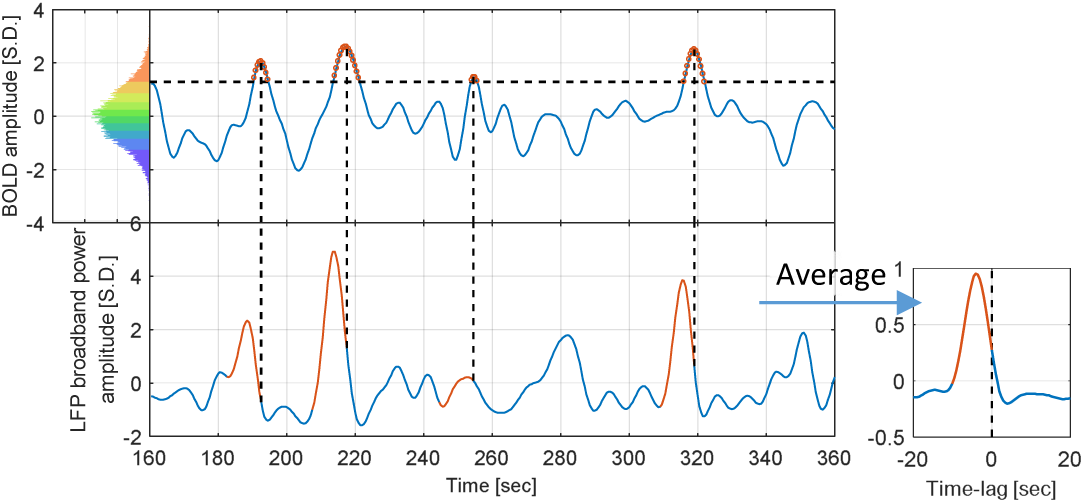
Illustration of the procedure to obtain the BOLD-triggered average time course. The data shown is from the ISO group (32 scans, each containing 1000 time points). The pooled distribution is evenly divided into 10 groups, each containing 3200 samples (red for high BOLD value and blue for low BOLD value, this color-coding applies to all figures that involve BOLD amplitude groups). A 200-second segment of a particular scan is plotted to illustrate the procedure. For a selected BOLD group (90∼100% BOLD is shown), the thresholds were obtained from the pooled distribution (>1.28 BOLD S.D. for this group), and applied to the time course to identify those time points as the BOLD events (marked in red circles on the top). Then the corresponding LFP segments proceeding the BOLD events were extracted (red segments on the bottom) and averaged across all 32 scans to obtain the BOLD-triggered average time course. For display purposes, only four LFP segments were shown.

Note that the process can be applied to the LFP power in other frequency bands as well, and the BOLD-triggered averages obtained in different frequency bands are directly comparable with each other, because they all share the same BOLD triggers and are therefore aligned along the time axis. Figure 2 shows the BOLD-triggered average of LFP power in six frequency bands (delta 1∼4Hz, theta 4∼8Hz, alpha 8∼12Hz, low frequency beta 12∼25Hz, high frequency beta 25∼40Hz and gamma 40∼100Hz) as well as the broadband power, which is the sum across the six frequency bands.

**Figure 2.**
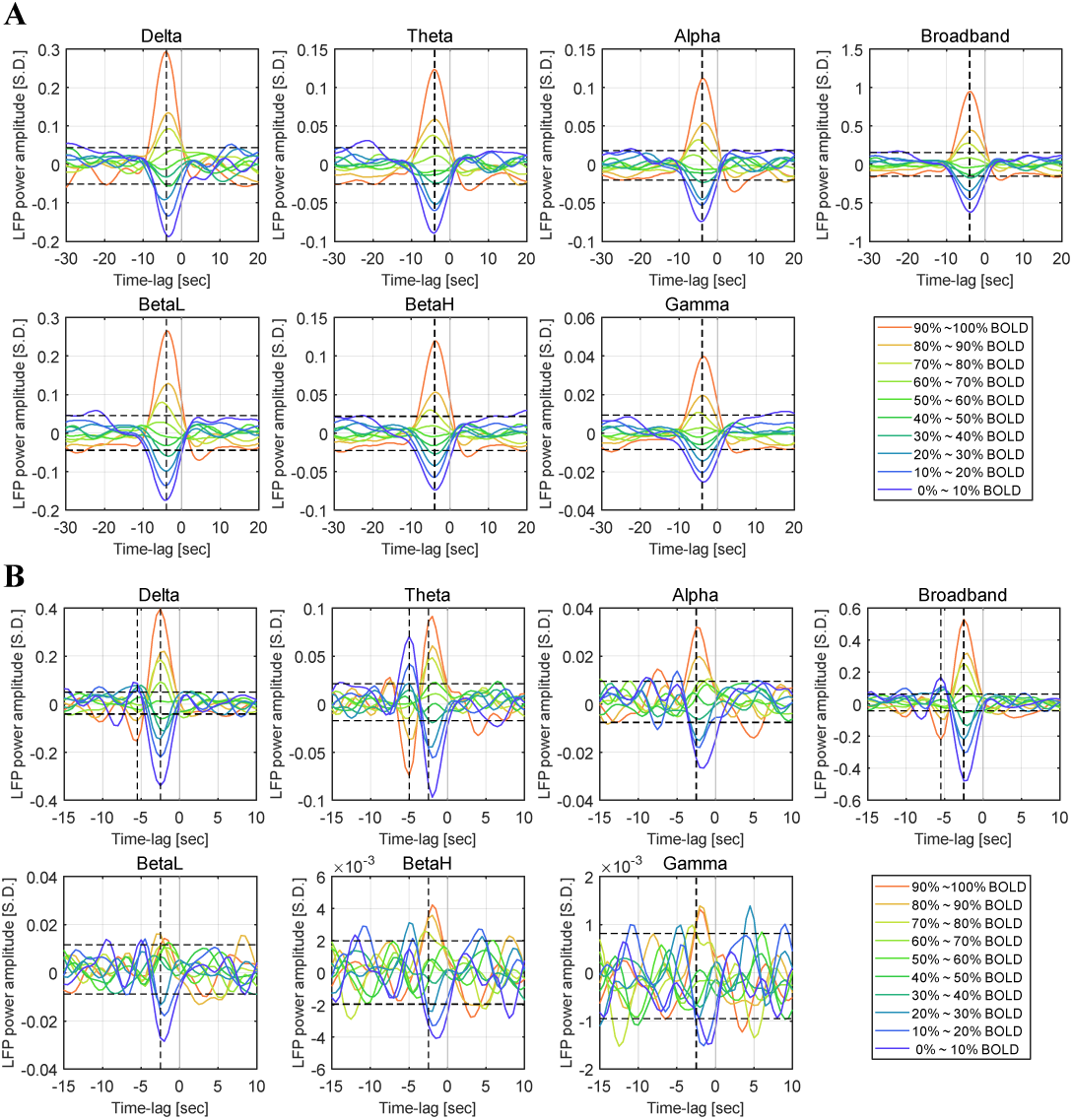
Time courses of the BOLD-triggered average of LFP power in different frequency bands under ISO anesthesia (panel A, 32 scans) and DMED anesthesia (panel B, 22 scans). The x-axis is the time lag with respect to the BOLD triggers. The y-axis is the normalized LFP power (the standard deviation of broadband LFP power is 1 LFP S.D.). The vertical line shows the maximally-correlated lag (−4 seconds for ISO in all frequency bands; −2.5 second for DMED in all frequency bands, with an additional line for the negative deviation which is located at −5.5 seconds for Delta band and Broadband, and at −5 seconds for Theta band). Each point in the time courses is the averaged value of roughly 10% of the dataset. The upper threshold and lower threshold for 95% confidence intervals are obtained from the 97.5 and 2.5 percentiles of the noise empirical distribution, respectively. BetaL, low frequency beta 12∼25Hz, BetaH, high frequency beta, 25∼40Hz.

Generally speaking, the BOLD-triggered average shows that the LFP-BOLD relationship is monotonic. Before the strong positive BOLD events (70∼100 percentiles), the LFP power exhibits an increased amplitude, whereas before the strong negative BOLD events (0∼40 percentiles), the LFP power exhibits a decreased amplitude. The peak change occurs at a time lag in line with previous findings (16), which is 4 seconds under ISO and 2.5 seconds under DMED. For the BOLD events with an amplitude around the median value (40∼70 percentiles), the LFP power shows a consistent trend (LFP power increases before a BOLD event higher than median value, and decreases before a BOLD event lower than median value), however, the effect is not significant enough to be distinguished from the general noise of the time courses. The 95% confidence interval of the random fluctuations is estimated by manually labeling the segments of BOLD-triggered average with a lag of −30 ∼ −15 seconds and 5 ∼ 20 seconds as the “noise” or irrelevant time segments, which provides 600 time points for the empirical estimation of the noise distribution. The aforementioned LFP power percentiles that significantly differ from the noise (70 ∼ 100% for positive BOLD events and 0 ∼ 40% for negative BOLD events) show some minor variations in the exact percentile values, depending on the anesthetic agents and LFP frequency bands.

Aside from the general trend, there are some additional patterns observed in DMED that are worth noting. First, as the frequency increases from low frequency beta band to gamma band, the time course become heavily contaminated by noise, and most of the percentile levels are no longer significantly different from the noise. This is potentially caused by the low signal-to-noise ratio (SNR) in these high frequency bands under DMED anesthesia, because the energy distribution of LFP decays much faster as frequency goes up when compared to ISO anesthesia (see Figure A.4). Secondly, there is a bipolar structure in delta and theta bands, which means that on average, in addition to the main peak at 2.5 seconds before the event, there is a secondary peak with inversed polarity at 5 seconds before the event, suggesting an anti-correlation between LFP power and BOLD at the time-lag of −5 seconds in these two bands. Since most of the energy is distributed in delta and theta bands, the broadband time course looks like a blend of these two bands.

Figure 3 shows the scatter plot of LFP power and BOLD at the maximally-correlated lag (−4 seconds and −2.5 seconds under ISO and DMED respectively), which provides a general idea about how LFP power is distributed in each BOLD level. It can be seen that while the average value within each cluster (white line plot) shows a clear correlation with BOLD amplitude, individual points have a very widespread distribution. Even for the highest 10% BOLD events, there are still a large number of LFP power occurrences that are below zero. So the increase of LFP power before a high BOLD event shown in Figure 3 is only an average effect, meaning that a high BOLD event does not guarantee an increase in the LFP power. The detailed distributions are provided in Figure A.3.

**Figure 3.**
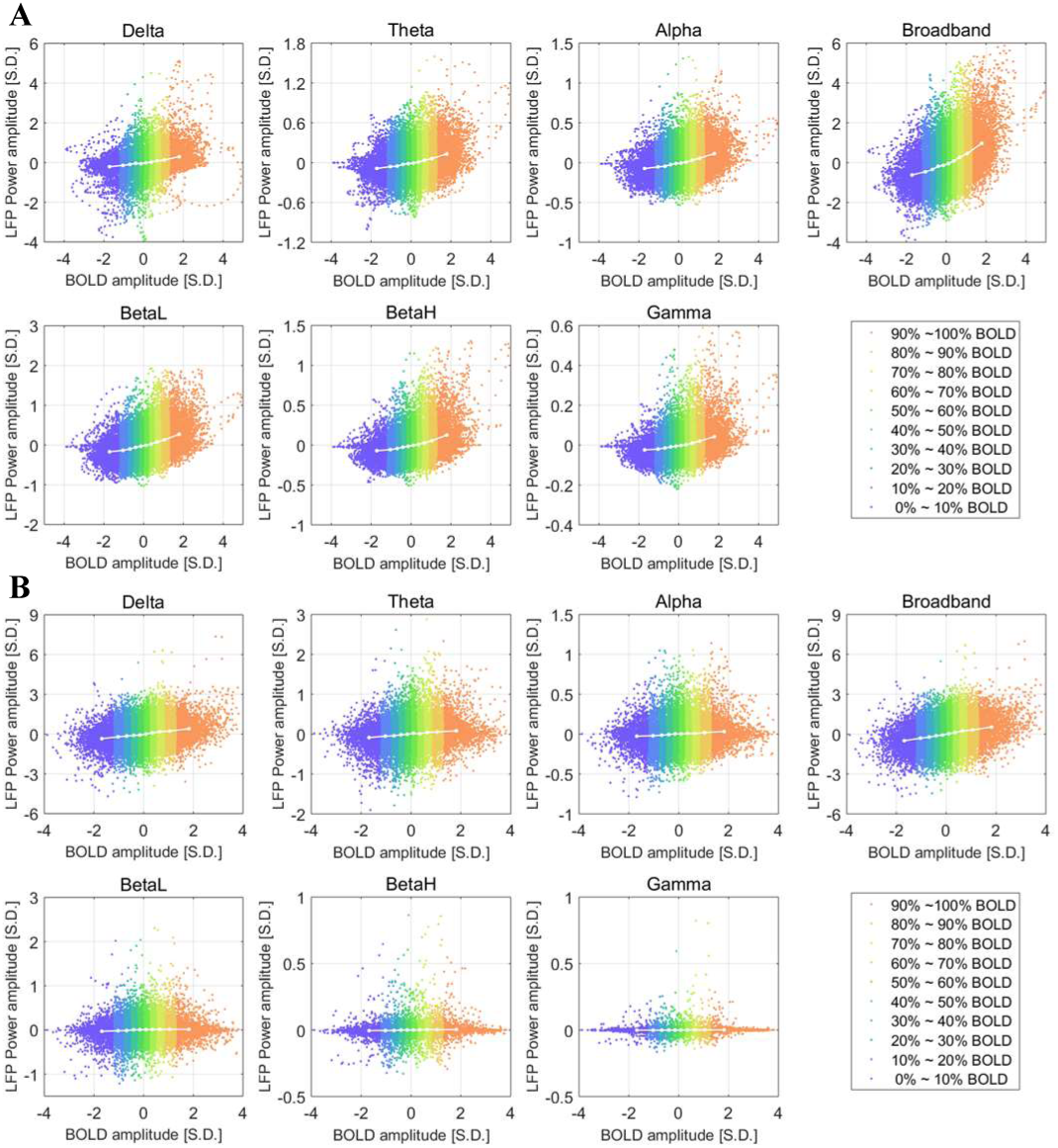
Scatter plot of LFP vs BOLD and the centroids of each BOLD group under ISO (panel A) and DMED (panel B). Each dot represents a time point with its BOLD value and LFP power value. The centroids are essentially showing the BOLD-triggered averages at the maximally-correlated lag, which are the value specified by the cursors in Figure 3.

### 2.2. LFP Co-activation and Co-deactivation Patterns

The BOLD-triggered average of LFP power time courses shows that only a portion (0∼40% and 70%∼100%) of the time courses can be distinguished from the general random fluctuations of the BOLD-triggered averages, which suggests that only LFP power higher than a certain threshold will trigger a BOLD response (in terms of the averaged effect). This finding is somewhat similar to the findings in co-activation patterns (CAPs) where it is revealed that the interregional BOLD correlations result from instantaneous co-activations of multiple brain regions at some critical time points rather than from continuous, sustained interregional neuronal interactions. We hypothesized that these two findings are tied together and the correlation between BOLD and LFP may be driven by instantaneous co-activations or co-deactivations as well, which might be the physiological reason why the interregional BOLD correlations are driven by instantaneous events.

To test the hypothesis, we proposed a modified version of CAPs that is obtained by applying thresholds to the LFP broadband power time course, as opposite to the original method, which applies thresholds to a selected seed region in the BOLD image series. In this article, the former method is referred to as LFP-CAPs and the latter one is referred to as BOLD-CAPs. To calculate LFP-CAPs, first several thresholds were calculated from the percentiles of LFP broadband power. For any given threshold, whenever LFP broadband power surpasses the threshold, the corresponding BOLD time frame (4 seconds under ISO, 2.5 seconds under DMED) succeeding the event was extracted. Each voxel was then averaged over the selected BOLD time frames, and the final averaged map was compared with the cross-correlation map between LFP broadband power and BOLD image series using the entire time course. In additional to LFP-CAPs, the original BOLD-CAPs were also calculated for comparison.

Figure 4 shows that as more and more frames are included by lowering the threshold, the spatial similarity between CAPs and the correlation maps increases rapidly, and reaches a plateau above 0.967 after including 10% of the data, suggesting the highest 10% LFP events or BOLD events can accurately replicate the spatial structure in correlation maps. Even a single frame is able to provide a general shape of the correlation map, which implies that the high amplitude events (either high LFP or high BOLD) are dominating the functional networks. These results are generally in line with Liu et al. 2013 (10).

**Figure 4.**
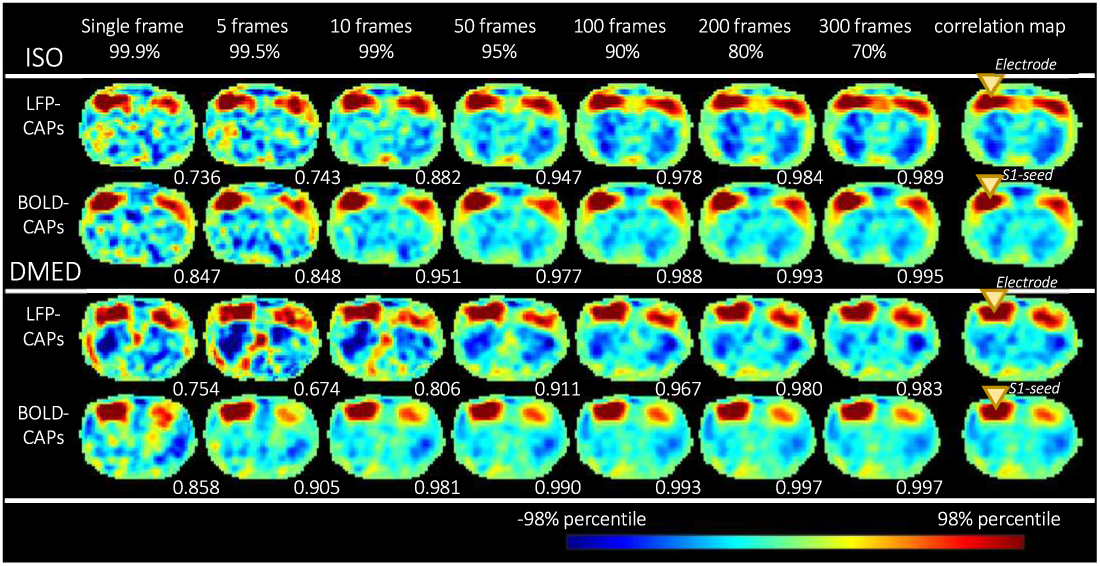
Co-activation patterns become more similar to the correlation map as more frames are included for calculation. From left to right, as the threshold (shown in percentiles) become lower, more frames are included. The LFP-BOLD correlation map (at the maximally correlated lag) and BOLD S1-seeded correlation map are shown on the far right (the yellow triangles show the location of the electrode and the S1-seed). For each CAP, the spatial similarity with regard to the correlation map is shown in the bottom right corner of each image. It can be seen that the spatial similarity increases very quickly as more frames are included, and reaches a plateau near 1 when a certain amount of frames are included. Even if only 10%∼20% of the dataset is used, most CAPs can replicate a spatial pattern nearly identical to the correlation map, which is calculated from the entire dataset. Each image is normalized by its 98 percentile to enable easier comparison of the spatial patterns.

In addition, the highest 15% BOLD time frames were selected for temporal decomposition (the BOLD-CAPs only need 5% to resemble the spatial pattern, but to avoid randomness caused by extremely small sample size, the threshold was set to 15%). K-means clustering was performed on the selected BOLD time frames (the distance was defined as 1 minus Pearson correlation coefficient), and the within cluster averages produce the temporal decomposition of the co-activation patterns. The number of clusters k was chosen to be 6 instead of 8 or 12 to avoid over interpretation, because the single slice images generally contain less spatial information compared to the 3D images shown in the original CAP paper.

It can be observed both visually from the patterns themselves and the similarity matrix that, despite some differences, most LFP-CAPs and BOLD-CAPs are highly spatially similar to each other (see Figure 5). The 2 by 2 quasi-diagonal elements in bright yellow can be easily distinguished from the other elements, suggesting that the intra-CAPs similarities are high among LFP-CAPs and BOLD-CAPs, and the inter-CAPs similarities vary. It is worth noting that the similarity between LFP-CAPs and BOLD-CAPs is not the result of overlapping triggers, because the high LFP events and high BOLD events only have 25.6% overlap in the time domain (see Figure A.5). Other factors (like network structure) that affect network dynamics may account for the similarity. There is also an intra-CAPs similarity observed across ISO and DMED (marked by the red dotted line), which can be distinguished from other inter-CAPs similarities, but the correlation values are not as high as the ones within the same anesthetic agents. This suggests when the anesthesia changes from ISO to DMED, the instantaneous functional networks change in some ways, but retains certain properties of spatial organization. It is worth noting that the correlation maps obtained from the conventional correlation analysis are visually very similar in these two cases despite the distinct mechanisms of anesthesia under the two agents. This suggests that the CAPs method is more sensitive to anesthetic-related changes than the conventional correlation analysis, possibly because it better preserves information about network dynamics.

**Figure 5.**
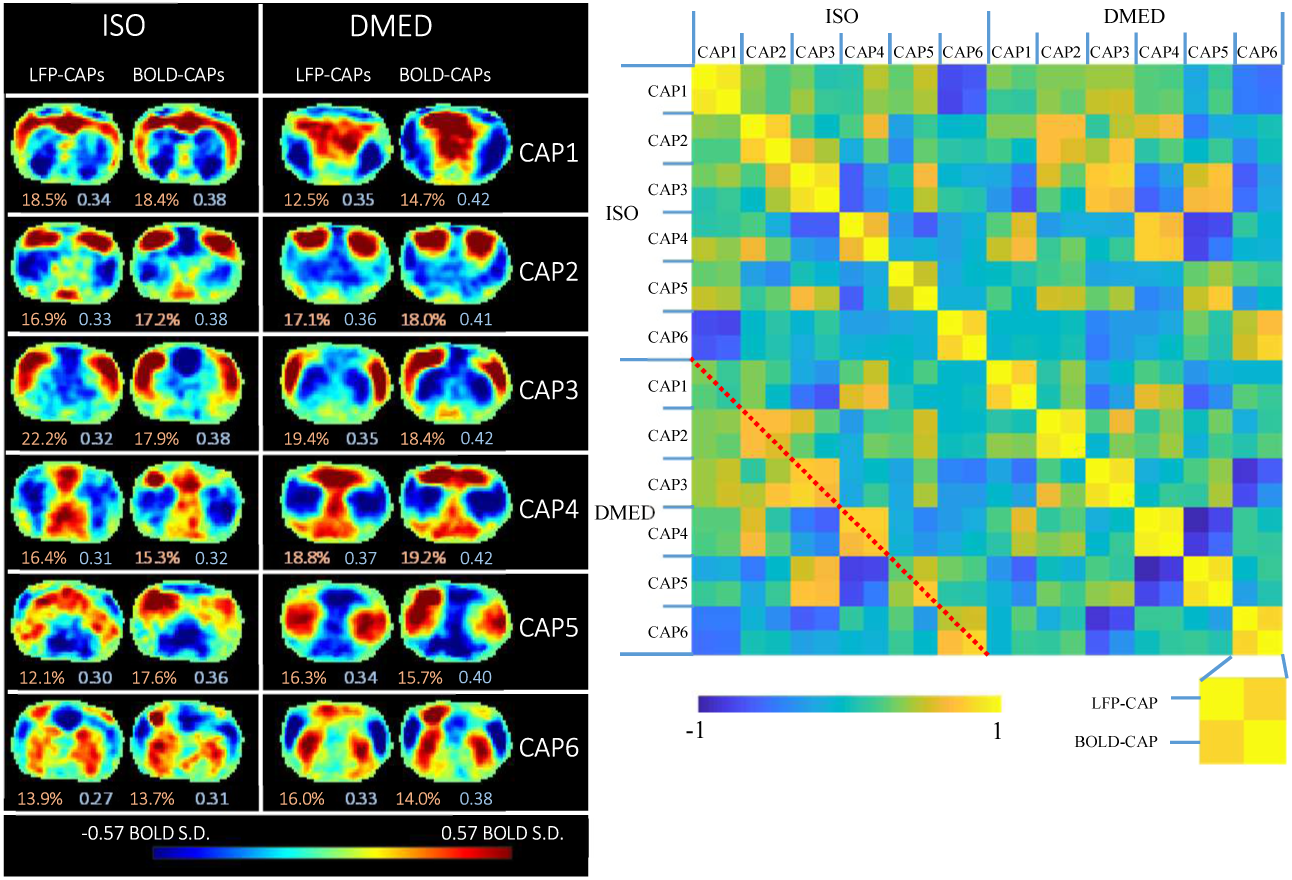
Temporal decomposition of the selected co-activation frames. The frames selected based on high amplitude events are further divided into six clusters using the k-means algorithm (k=6) to produce CAPs. The threshold used for selecting frames was 15% for both LFP-CAPs and BOLD-CAPs. The LFP-CAPs under ISO are sorted based on the consistency (the average spatial similarity of each fMRI frame to the group mean). The LFP-CAPs under DMED are sorted to maximize the summed spatial similarity between LFP-CAPs under ISO and LFP-CAPs under DMED (for easier comparison across different anesthesias). The BOLD-CAPs are also sorted in a similar way using LFP-CAPs as the benchmark. The consistency (light red) and fraction values (light blue) of the CAPs are shown on the bottom of each image.

## 3. Discussion

### 3.1. General Findings

In this study, we utilized two complementary methods for the analysis of simultaneous rs-fMRI and LFP recording data to show that the correlation between BOLD and LFP is mainly influenced by discrete high amplitude events. The high amplitude LFP power events not only show an increased likelihood of eliciting a localized BOLD response, but also produce co-activation patterns that are nearly identical to the cross correlation map between LFP and BOLD. Furthermore, the co-activation events can be clustered into several distinct groups, providing electrophysiological evidence of neural underpinnings for the multiple BOLD-CAPs obtained from rs-fMRI data.

### 3.2. On average, the LFP power preceding the BOLD events exhibits a stereotypical profile

BOLD-triggered averaging revealed that there are certain temporal patterns that manifest before a BOLD event. The increase or decrease of LFP power from baseline peaks at 4 seconds before the BOLD response under ISO, and 2.5 seconds under DMED. This finding is in agreement with the previous studies using conventional cross correlation, but provides more information about the behaviors of subsets of BOLD time points that have different amplitudes. On average, the high amplitude positive BOLD events (70∼100 percentiles) are preferentially preceded by an increase in LFP power, and the negative BOLD events (0∼40 percentiles) prefer a decrease in LFP power. This amplitude preference vanishes when the amplitude of BOLD falls into the range near median value (40∼70 percentiles), because the changes in LFP power are no longer distinguishable from the random fluctuations. The fact that about 30% of BOLD time points near the median value have no consistent correspondence with the LFP power implies an intrinsic limitation in extracting function connectivity patterns using cross correlation, because a great amount of data that contains little information about neural activity is influencing the result.

Note too that decreases in LFP power precede decreases in the BOLD signal. The origin of the negative BOLD signal has long been debated, given its complicated dependence on neural and vascular parameters. Here we show that during spontaneous activity, decreases in LFP power are closely linked to decreases in the BOLD signal. While this does not rule out other contributions to the negative BOLD effect seen in fMRI (e.g., vascular steal), it does provide support for a neural basis as well.

### 3.3. The correlation between LFP and BOLD is driven by a few distinct events

The results of LFP-CAPs suggest that, selecting the fMRI frames with the highest 15% of LFP amplitudes can accurately reproduce the spatial patterns seen in the cross correlation map, which requires utilizing the full dataset. This together with the thresholds found in BOLD-triggered average analyses suggest that the relationship between LFP and BOLD is dominated by a few distinct events, specifically those with LFP power amplitude higher or lower than a certain threshold. This finding in a way matches with the previous studies performed by Liu et al. (10) and Tagliazucchi et al. (11), where it is revealed that the interregional correlation in fMRI are driven by instantaneous BOLD events. Our work confirms that the same principle holds for LFP power recorded at a particular location and the BOLD signal, such that the relationship between the two is driven by a few high amplitude events.

The scatter plots of BOLD vs LFP power make it clear that despite the average relationship between high amplitude LFP events and high amplitude BOLD events, the relationship is highly variable and strong BOLD events can actually be tied to decreases in LFP power. If it were possible to identify the points where BOLD was linked to LFP power (e.g., by incorporating additional information into the analysis), the interpretation of the BOLD signal could be made easier. Unfortunately, our preliminary analysis that considered the length of the LFP burst along with its amplitude did not improve our ability to determine which strong BOLD responses reflected high amplitude LFP events. It may be the case that the noisy relationship between the two signals is fundamental to the processes that mediate neurovascular coupling.

### 3.4. Frequency Dependence of the BOLD-LFP Relationship and Other Findings

It was reported previously that the BOLD signal preferentially correlates with specific frequency bands of the LFP. The frequency ranges reported were gamma band under ISO and fentanyl (9, 13), gamma band under fentanyl and thiopental (17), alpha to gamma band under remifentanil (14), delta to gamma band under ISO (12, 16), and delta and theta band under DMED (16). The BOLD-triggered average presented in this study is consistent with a frequency preference that is dependent on the anesthetic agent. There is no significant difference among different frequency ranges under ISO, while under DMED, the BOLD-triggered average appears noisier as the frequency increases, especially in the gamma band. These findings are in agreement with the correlation and coherence analysis reported in Pan et al. (16), partly because the two studies share similar data preprocessing pipeline and some common datasets. However, other researchers conclude that oftentimes gamma band is the most informative band (9, 13, 14, 16). The differences might be the result of a shift in the LFP power distribution towards lower frequencies under anesthetics like DMED, which results in very little power at the higher frequencies. This suggests that broadband power, rather than power in any particular band, may prove more a robust predictor of the BOLD signal across anesthetic protocols.

Aside from this, we also found an unexpected phenomenon, which is the bipolar structure in delta and theta bands under DMED. The secondary negative lobe peaks at −5 seconds, which suggests a strong anti-correlation exists between LFP and BOLD. However, this can be only interpreted as an averaged effect obtained from retrospectively regrouping the BOLD time points and may not be essential to trigger the BOLD events. We suspect it arises from the enhancement of delta and theta activity under DMED, creating ongoing oscillations in these frequencies.

Furthermore, a nonlinear relationship between LFP and BOLD is observed under ISO but not DMED. This can be seen from Figure 2 panel A, where the increase of LFP in positive BOLD percentile groups generally has a larger magnitude than the decrease of LFP in negative BOLD groups, and Figure 3 panel A, where the LFP-BOLD line plot clearly shows a curvature. This is a direct evidence that the relationship between LFP and BOLD can be nonlinear and depends on the anesthetic agents. Given the presence of nonlinearity, analysis methods that do not assume linear dependency should be considered instead of Pearson correlation (e.g., mutual information methods presented in Magri et al. (14)).

### 3.5. Methodological Comparison

In Magri et al. (14), the high gamma LFP power was used as the trigger and the BOLD time courses after the LFP events were averaged. The triggers were then subdivided into three groups based on alpha band or beta band amplitude. While this method is able to show that alpha and beta bands contain complementary information to gamma band, it is difficult to analyze the other frequency bands in any given subgroup, because there are too many possible combinations of the subgroups in different frequency bands. The mutual information analysis was performed prior to this LFP triggered BOLD averaging process to find the most informative frequency bands, (in their study gamma, beta and alpha were the three most informative ones), and avoid the enumeration of all possible combinations of different frequency bands. Another way to address the problem is to use BOLD as the trigger instead, as in this study. In the BOLD-triggered average, all of the frequency bands share the same BOLD triggers, so they are directly comparable.

Generally speaking, when compared to conventional analysis methods, the methods presented in this study have several advantages. First, they do not assume a linear dependency like correlation analysis does, and the BOLD triggered average was able to discover a nonlinear relationship. Secondly, they emphasize individual events, instead of continuous interactions. This enables both methods to provide more details about the dynamic aspects of the signal.

### 3.6. Implication for Future Studies

Time courses from rs-fMRI are often treated as continuous measurements of the hemodynamic response to neural activity. In reality, most of the information in the time courses appears to be carried by a much smaller set of discrete events, which in turn reflect discrete LFP power events. The finding that only strong LFP events result in a detectable hemodynamic response places fundamental limits on our ability to monitor brain dynamics with hemodynamic-based methods. While low-amplitude ongoing activity in the brain is critical for normal function, only strong excursions from the baseline activity will be detected. This suggests that rs-fMRI should be modeled as a discrete series of states rather than a more continuous evolution over time.

The thresholds for separating events from background fluctuations are dependent on the number of subjects as well as the duration of each scan. By including more data, one can reduce the random fluctuations and thus make the temporal structure of the BOLD-triggered average clearer (similar to SNR). In our analysis, we included 32 scans under ISO and 22 scans under DMED, each lasts 8 min 20 sec. For any single scan without extended acquisition time, the amount of irrelevant data could be more than 30%.

### 3.7. Technical Limitations

The level of ISO impacts neuronal activity and cerebral perfusion. Mixing datasets with different ISO levels, as done here, could potentially introduce variability in either the BOLD-triggered average time courses or the LFP-CAPs. However, it should be noted that all ISO rats were imaged while in the burst-suppression regime. The number of scans was too small to perform further separation by ISO level. In the future, the impact of varying ISO level on the BOLD-triggered average is worth investigating, and more datasets under different ISO levels are needed.

The datasets used were deliberately chosen to have high correlation at the lags corresponding to previously observed hemodynamic delays (−4 seconds under ISO and −2.5 seconds under DMED). While this ensures the overall quality of data, it may introduce some bias as well. In all of our studies, we have found that the correlation coefficient drops drastically in some of the scans even when other scans of the same rat under the same anesthesia with almost identical physiological condition show high correlation. Minor fluctuations in temperature, slow changes in respiration, or other physiological effects of the anesthesia may result in poor data for one scan, which then returns to normal as the physiological parameters are corrected. We monitor rats undergoing simultaneous LFP and MRI very carefully, but it is difficult to keep their condition absolutely stable. The data used in this study were selected conservatively to ensure optimal results.

The LFP-CAPs utilize the spatial information obtained from fMRI, taking advantage of the fact the fMRI data are more densely sampled in spatial domain than LFPs, providing insight into time-varying activity that could not be discovered using methods like correlation analysis. However, this approach is still not ideal, because although LFP is a more direct measurement of the neuronal activity, the LFP appears to be more loosely connected with BOLD signals in other regions, when compared to the BOLD seeded drawn near the electrode (because LFP-CAPs need more frames to reproduce the spatial pattern, as shown in Figure 4). So far we can only confirm that the LFP co-activates with BOLD signals in a similar way that the BOLD seed does, and such similarity can only imply that the apparent time-varying functional connectivity observed in CAPs may be the result of the co-activations of neurons. To collect direct evidence that the resting state networks are dynamic and are driven by discrete events, one needs to use multiple electrodes located in areas throughout the network. Such an experiment has many technical difficulties, and LFP-CAPs remain a good alternative to provide spatial information until the successful implementation of multi-region, high-resolution LFP recording and MRI.

## 4. Conclusion

To conclude, in this article we proposed two methods to analyze simultaneous fMRI and LFP recording data: the BOLD-triggered average and the LFP-CAPs. The BOLD-triggered average shows that the there is a particular temporal pattern in the LFP power shortly before any type of BOLD events, especially those events with the highest BOLD amplitudes where the pattern of LFP power stimulus is most easily distinguished from the background noise. Under different anesthesia, the temporal patterns in the averaged LFP power show different frequency preferences.

The spatial similarities between the LFP-CAP and the cross correlation map suggests that the relationship between LFP and BOLD is driven by instantaneous co-activations or co-deactivations, which is in line with the finding in BOLD-triggered averages that the averaged LFP stimulus will exhibit some noticeable patterns only if the BOLD triggers are high in amplitude The spatial similarities between LFP-CAPs and BOLD-CAPs suggests that the time-varying resting state networks found in fMRI studies may be attributed to the time-varying behavior of LFP in different brain regions, although the underlying mechanism is still not fully understood.

## 5. Materials and Methods

### 5.1. Animal Preparation

All animal experiments were performed in compliance with NIH guidelines and were approved by the Emory University Institutional Animal Care and Use Committee. Previously acquired data from 36 Sprague–Dawley rats (male, 200–300 g, Charles River) were used in this study. A full description of the methods is given in the prior publication (**Error! Reference source not found**.2) and summarized here. All rats were anesthetized with 2% isoflurane during surgery. Fine tip electrodes (∼10 μm in diameter, borosilicate pipettes) were prepared with micropipette pullers (PE-2; NARISHIGE). The electrodes were filled with artificial cerebrospinal fluid (ACSF), resulting in an impedance of 1 – 5MΩ between the chloridized silver wire and the extracellular environment. The details of the surgical procedures and microelectrode implantation have been described in (18). A pair of micro-glass electrodes was implanted in S1FL of the left and right hemispheres separately and secured to the skull using dental cement (methyl methacrylate) for all rats. In order to reduce the MRI artifacts caused by susceptibility, a layer of toothpaste (Colgate, NY) was applied to replace the removed skin and muscle over the skull, and also the dental cement was used at the area >0.5 mm posterior to the imaging slice.

### 5.2. Simultaneous fMRI imaging and LFP recording

All imaging was performed on a 9.4T/20 cm horizontal bore small animal MRI system (Bruker, Billerica, MA). A three-plane scout image was first acquired to position the fMRI scans. To improve the homogeneity of the magnetic field, the volume of interest (6 mm^3^) was shimmed using FASTMAP (19). Manual shimming adjustment was then applied when necessary to improve the field homogeneity of the selected slice. For fMRI studies, a coronal imaging slice was selected to cover bilateral S1FL areas, in which the glass recording electrode tips were implanted. The EPI imaging parameters were FOV, 1.92 × 1.92 cm^2^; matrix size, 64 × 64; in-plane resolution, 0.3 × 0.3 mm^2^; slice thickness, 2 mm; and TR/TE, 500/15 ms. A total of 337 resting state scans were collected under several different concentrations of isoflurane ranging from 1% to 2%. (ISO, n = 100) or dexmedetomidine (DMED, n = 237) anesthesia. The isoflurane, in a mixture of 70% O2 and 30% room air, was continuously delivered to the nosecone, allowing for free breathing throughout the experiment. The rat’s oxygen saturation, measured with a pulse oximeter, was kept above 98% throughout the data acquisition process. For DMED studies, a bolus of 0.025 mg/kg dexmedetomidine was injected subcutaneously. Isoflurane was disconnected 10 min afterwards, and switched to a continuous subcutaneous infusion of dexmedetomidine (0.05 mg/kg/h). The dose was increased by a factor of three (0.15 mg/kg/h) after ∼1.5 h, following the protocol for prolonged sedation described in (20). The DMED scans were conducted >3 h after switching from ISO to avoid any residual ISO effects (21). The fMRI image acquisition lasts 8 min 20 sec (1000 TR), and 20 dummy scans were acquired to reduce transient signal intensity fluctuations at the start of the image series, which makes the total length of LFP segments that contains gradient-induced artifacts become 8 min 30 sec. Because the whole dataset was acquired over a period of several years, there were two sets of recording parameters: 1. (×500 amplified, 0–100 Hz bandpass-filtered, 60 Hz notch-filtered, 12 kHz sampling rate, and ∼10 min acquisition length) and 2. (×1000 amplified, 0.1 Hz–5 kHz bandpass-filtered, 60 Hz notch-filtered, and 12 kHz sampling rate, and ∼14 min acquisition length). The LFP recording lasts longer than the image acquisition to record the LFP without the gradient-induced artifacts, which provides a benchmark for the artifact-removal algorithm. There are two LFP recording segments before and after the image acquisition that last either around 1 min or around 3 min, depending on the parameter sets. However, these differences in the recording parameters will not compromise the analysis because in the data processing the LFP was all band-pass filtered to 0.1-100Hz, the amplitude was normalized, and the excessive LFP segments were truncated to match the length of fMRI data. A 16 bit analog-to-digital converter (PCI-6281; National Instruments) was used for analog to digital conversion. All physiological parameters were monitored and maintained within normal ranges, including rectal temperature, respiration rate, SpO2/cardiac rate. The animals were sacrificed at the end of the experiment.

### 5.3. LFP Data Pre-processing

The electrophysiological signal recorded during fMRI scans contains neural signals and gradient-induced artifacts, which are easily distinguished by their high amplitudes. These gradient-induced artifacts were removed offline in MATLAB (Mathworks) following a similar procedure to that illustrated in (12). To identify the time when gradient-induced artifacts are present, the rising edge is captured by comparing the first order derivative with a predefined threshold. If the total number of the detected gradient-induced artifacts equals to 1020 (the duration of fMRI scan was 1000 TRs, and there were 20 dummy scans before the actual data acquisition), the LFP data was then proceeded to the next step. Otherwise if the artifacts are not identified correctly due to the presence of some large noise spikes, we first tried to manually replace those spikes with linear interpolations. If the artifact identification was still problematic after manual denoising, the dataset was discarded.

The LFP data can be divided into several segments using the triggers, and there should be 1020 segments that contains gradient-induced artifacts in each scan. Those segments were averaged to obtain the noise template, which was then subtracted from the individual segments of the raw LFP signal to get the denoised LFP signal. This method takes care of the gradient artifacts in the second phase (25 ms after the trigger), but the residual artifacts in the first phase (0∼25 ms after the trigger) were still overwhelming after subtracting the noise template, so the first 25 ms of the LFP data were replaced with linear interpolation.

The denoised LFP signal was then low pass filtered to 100 Hz using to remove any residual artifacts, and downsampled from 12KHz to 500Hz to reduce file size and computation cost. These downsampled LFP signals were then used to calculate the band-limited power (BLP) in different frequency ranges (Delta 1∼4Hz, Theta 4∼8Hz, Alpha 8∼12Hz, Beta Low 12∼25Hz, Beta High 25∼40Hz, Gamma 40∼100Hz) using sliding window. The sliding window has a length of 1 second and 50% overlap (so the window moves 0.5 second every time to match the temporal resolution of fMRI data, which is 0.5 second as well) and is centered at the echo time of the fMRI scan for each TR index. Within each window, the power spectral density (PSD) function was calculated using Welch’s method (4 segments, 50% overlap) and was integrated over different frequency bands to obtain the BLP time courses at the corresponding TR index. The BLP time courses were then band-pass filtered (0.01-0.1Hz for ISO and 0.01-0.25Hz for DMED (16)). After the pre-processing, the raw LFP data with a sampling rate of 12 KHz was converted to BLP time courses with a temporal resolution of 0.5 seconds and a duration of 1000 TRs, which is 8 min 20 sec long. For further BOLD triggered average analysis, the BLP time courses in every frequency band from the same scan was normalized by a common scaling factor such that the standard deviation of the broadband power is equal to 1, which makes the datasets with various amplitudes comparable with each other.

### 5.4. FMRI Data Pre-processing

The fMRI data was preprocessed in the following procedure. A brain mask for each scan was obtained from the first image of the scan using active contour methods and was dilated by 2 voxels before running motion correction on SPM 12. The motion corrected image series were then spatially smoothed using a Gaussian kernel with a FWHM of 2.8 voxel (2.8*0.3mm = 0.84mm). Then global signal and linear drift regression, as well as band-pass filtering (0.01-0.1Hz for ISO and 0.01-0.25Hz for DMED) were performed voxel-wise. As a quality assurance step, the cross-correlation map of LFP bandlimited-power versus BOLD is calculated at the lag when the correlation is expected to reach the maximum (4 seconds for ISO and 2.5 seconds for DMED). If there are high LFP broadband power versus BOLD S1-seed cross-correlation, and a bilateral symmetry in BOLD S1-seeded correlation map, the dataset was considered as high-quality data and was proceeded to the next step. By these criteria, 22 scans under DMED from 10 rats and 32 scans under various ISO concentrations from 12 rats were selected out of 337 scans from 36 rats. The selected datasets were then normalized voxel-wise to produce the BOLD image series for the co-activation patterns analysis. For BOLD triggered average analysis, a ROI was manually selected based on the cross-correlation map, and the BOLD signal was averaged over the ROI. Finally, the BOLD signal averaged over the ROI was z-scored so that the averaged BOLD signal is comparable with other fMRI scans.

### 5.5. Co-deactivation Patterns (CDAPs)

In addition to co-activation patterns, we also calculated co-deactivation patterns (CDAPs). The CDAPs were obtained by applying thresholds to the LFP broadband power time course, similar to calculating CAPs except the BOLD frames were selected when LFP broadband power was lower than the threshold, and the sign of the averaged intensity was flipped for better comparison with the correlation map. The comparison between CAPs and CDAPs are shown in Figure A.6 and Figure A.7. Although the CAPs described in the original paper lost similarities when the thresholds are too low (because the co-deactivation frames cancel out the effect of the co-activation frames), however, we found that those co-deactivation frames do not always destroy the patterns, and solely using them can produce a nearly identical spatial map, though they generally requires more frames to achieve the same level of similarities. So the fact that CAPs can reproduce correlation map should be interpreted as co-activation frames contain information that is sufficient, but not necessary to resemble the correlation map. The underlying reason might be the complicated coupling in the timing of CAPs and CDAPs, which is probably induced by the nature of temporal filtering with a relatively narrow frequency band of the power time course.

Another thing to notice is that, the amount of frames needed to reach the similarity plateau is different for different CAPs. Generally speaking, BOLD-CAPs and BOLD-CDAPs are the two fastest to reach the plateau (need 5∼10% of the data), followed by LFP-CAPs (need around 20%), and the LFP-CDAPs is the slowest (may need up to 30%). It is implied that since the spatial patterns in the correlation map can be accurately replicated by only a small portion of dataset, those selected frames contains all the information needed for resembling the pattern. So it is reasonable to assume that, if fewer frames is needed for resembling the pattern, then each frame contains more information than otherwise. Since LFP is linked to BOLD signal through intermediate steps including hemodynamic functions, whereas BOLD signal is more directly tied to the BOLD signals in other region through the same mechanism, it is not surprising to see that BOLD-CAPs reach plateau quicker than LFP-CAPs. The fact that LFP-CAPs need fewer frames than LFP-CDAPs implies that the LFP activation is more dominant than deactivation when driving the LFP-BOLD relationship.

## Acknowledgements

Funding sources: NIH 1 R01NS078095-01, BRAIN initiative and NSF INSPIRE. The authors would like to thank Chinese Scholarship Council (CSC) for financial support.

## Appendix

**Figure A.1.**
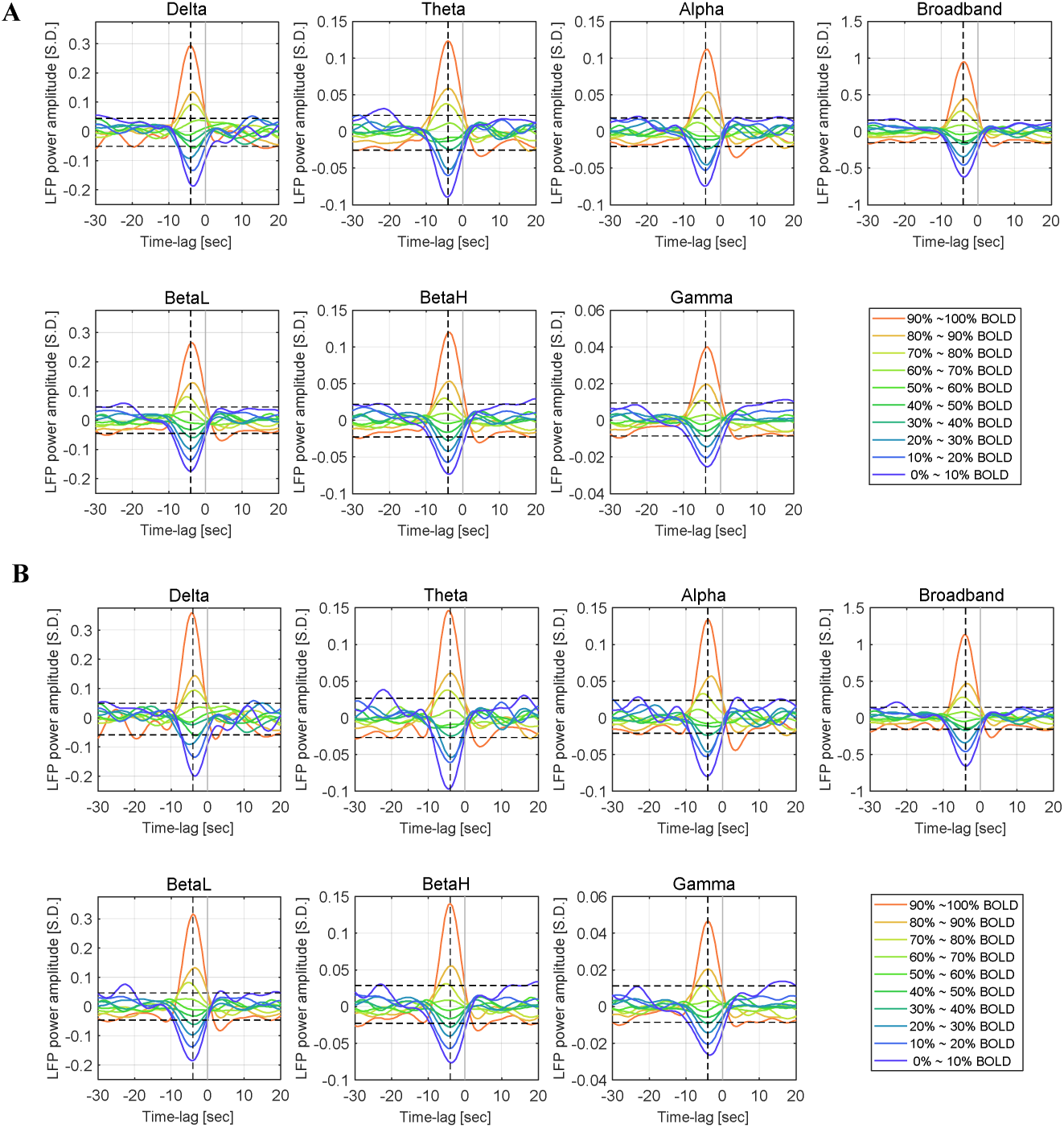
Two different strategies to get BOLD triggered average yield very similar results. Panel A, the BOLD triggered average obtained when treating each individual time point as a separated trigger, which is identical to the figure 2A except the scale of y-axis. Panel B, the BOLD triggered average obtained by combining adjacent triggers into one single trigger, weighted by the duration of the event. The two methods produce very similar results, the general shapes are nearly identical despite the later method gives higher amplitudes.

**Figure A.2.**
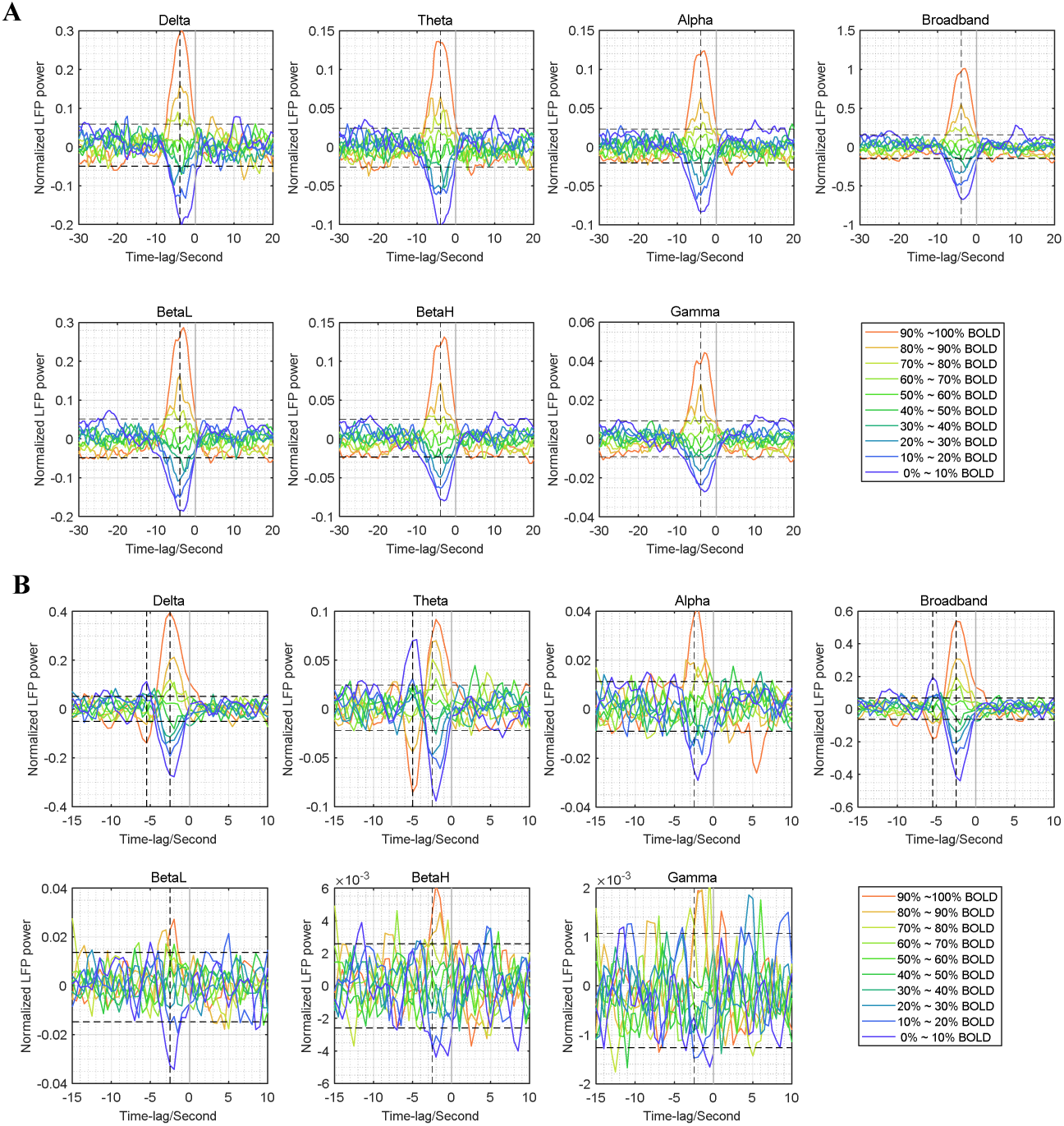
Unfiltered BOLD triggered average under ISO anesthesia (panel A) and DMED anesthesia (panel B). The BOLD time course for defining the triggers was still band-pass filtered (0.01∼0.1Hz for ISO and 0.01∼0.25Hz for DMED). Unlike figure 2, the LFP power time courses being averaged were unfiltered. The time course was sampled with a temporal resolution of 0.5 second, meaning any frequency components lower than 1Hz can be seen in the figure. However, despite of some minor shape distortion and noise contamination, the BOLD-triggered average time course is not heavily influenced by removing the band-pass filtering.

**Figure A.3.**
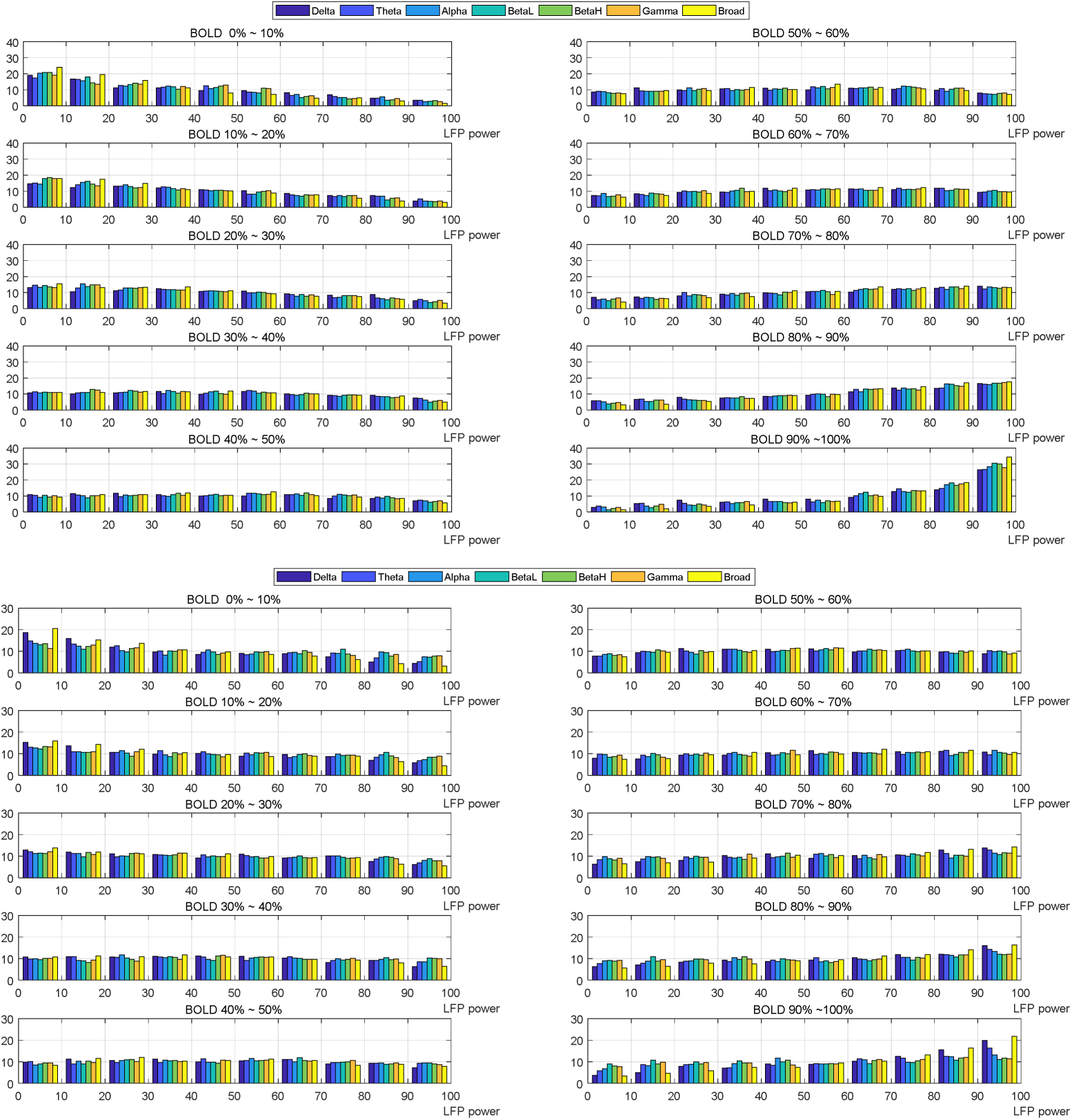
Histograms of LFP power under ISO anesthesia (panel A) and DMED anesthesia (panel B) at the maximally-correlated lag given a BOLD amplitude. The different colors shows different frequency component of LFP power. Each subplot shows that, given the BOLD amplitude is within in a certain range, the relative frequency in percentages (y-axis) of having a LFP power in a particular range specified by LFP power percentiles (x-axis). Everything is measured in percentiles so that the frequency bands with different amplitudes can be comparable. For any LFP power frequency band, the sum across 10 LFP power groups within a given BOLD group or the sum across 10 BOLD groups within a given LFP group is equal to 100%. It can be seen that under many circumstances (either 30∼70% BOLD or 30∼70% LFP) the relative frequency is around 10%, meaning it is almost uniformly distributed and thus knowing one signal does not help predicting the other. Only in the highest or the lowest BOLD groups, the distribution of LFP becomes skewed and is significantly different from a uniform distribution, and vice versa. This skewed distribution gives rise to the BOLD-triggered averages seen in figure 2 and motivated us to hypothesize that the correlation between LFP power and BOLD is driven by the most prominent events (either the LFP surpassing-threshold events or the BOLD surpassing-threshold events).

**Figure A.4.**
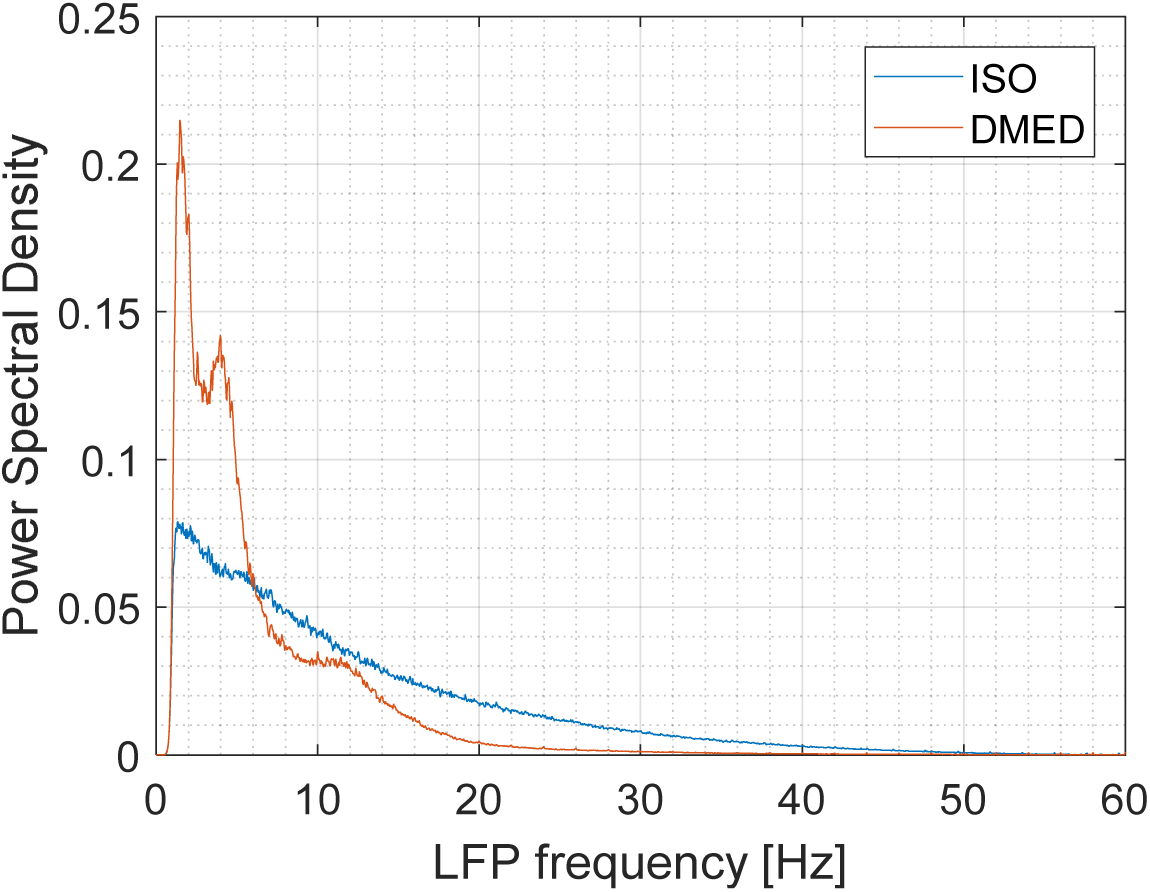
LFP power distribution (ISO, n=32, DMED, n=22). Each scan session (500 sec) was z-scored and the power spectral density (PSD) function was estimated using Welch’s method (50% overlap, 64 segments). Then the PSDs were averaged within ISO group and DMED group. The amount of energy distributed in delta, theta, alpha, low frequency beta, high frequency beta, gamma are 21.1%, 22.7%, 16.5%, 27.2%, 9.7% and 2.3% under ISO, respectively; and 44.4%, 28.7%, 12.9%, 11.9%, 1.3% and 0.4% under DMED, respectively. The PSD decays much faster in DMED as frequency goes higher, and the low energy in these high frequency bands makes the estimation of LFP power and therefore the BOLD-triggered average very noisy.

**Figure A.5.**
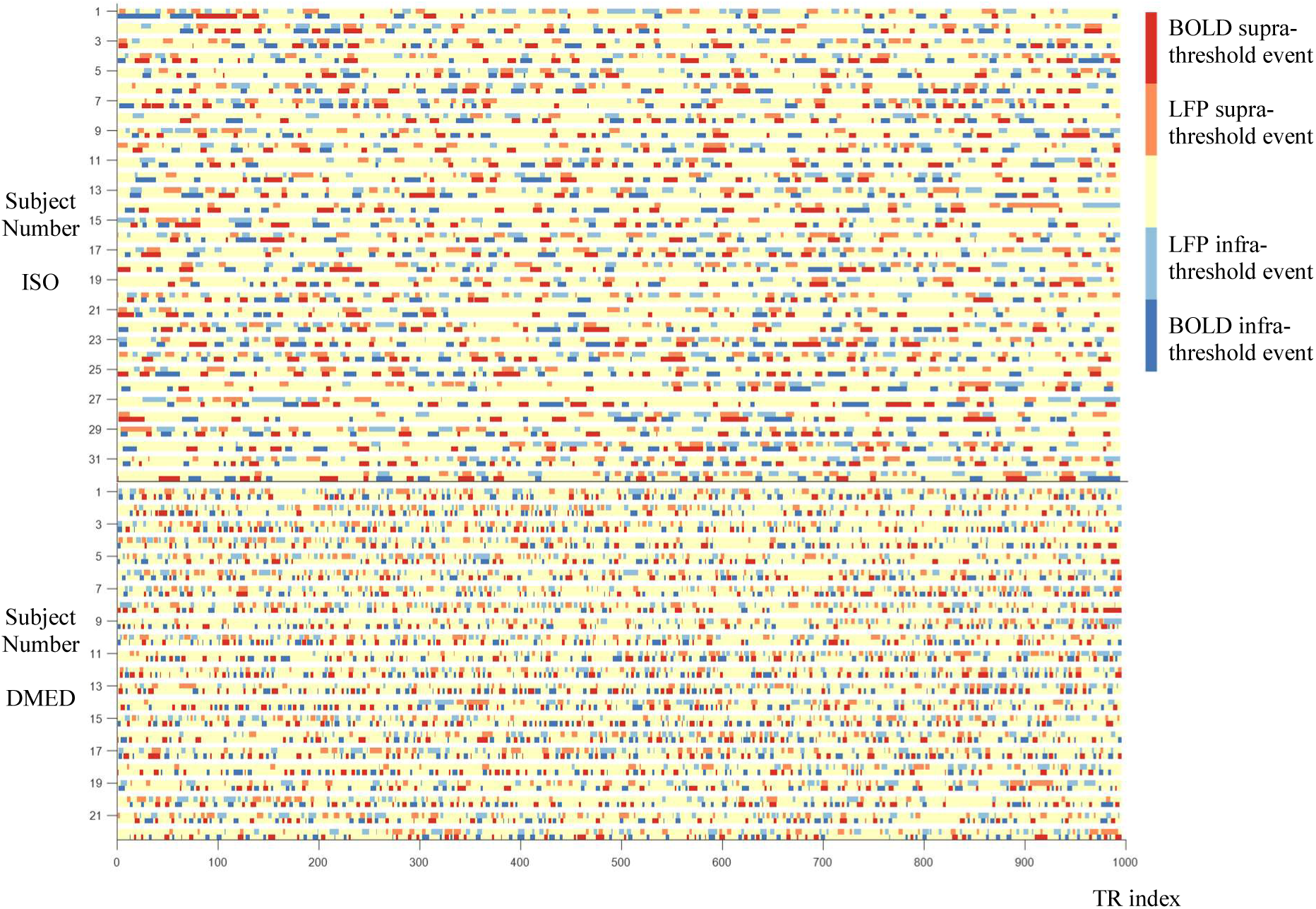
Timing of LFP and BOLD supra-threshold and infra-threshold events. The threshold was set to 15%. It can be seen that although sometimes the LFP and BOLD activate or deactivate concurrently, producing a positive correlation between the two, most of the time the LFP activations and BOLD activations do not overlap in the time domain (the fraction that overlap is 25.6% for activations and 24.3% for deactivations). Therefore the similarity between LFP-CAPs and BOLD-CAPs observed in Figure 5 might be attributable to other factors (like network structure) that might affect network dynamics, rather than the simple overlap in the timing.

**Figure A.6.**
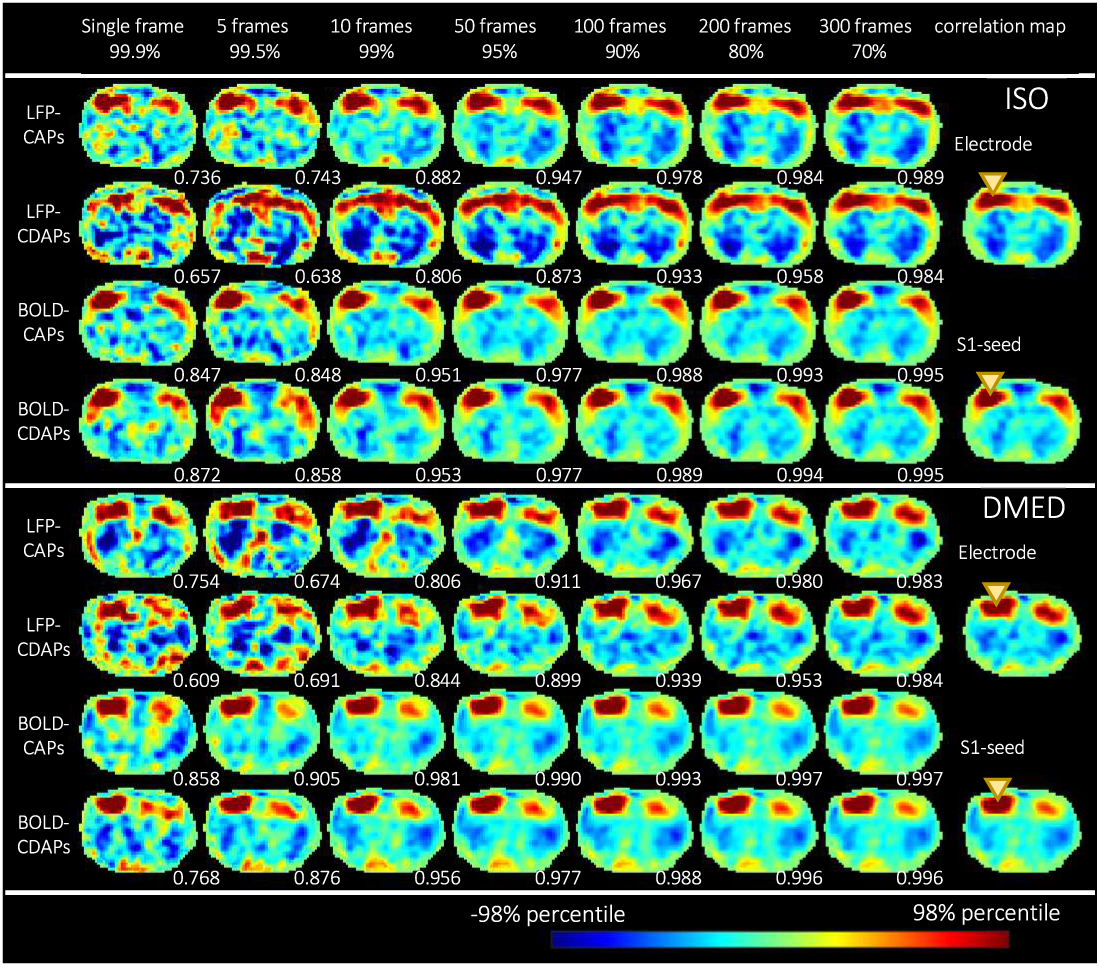
Co-activation patterns become more similar to the correlation map as more frames are included for calculation. From left to right, as the threshold (shown in percentiles) become lower (for CAPs, it would be the opposite for CDAPs), more frames are included (The number of frames shown specifies how many frames from each scan are selected for calculation, the total number of frames for ISO and DMED should be this number times the numbers of scans, which are 32 and 22 respectively). The LFP-BOLD correlation map (at the maximally correlated lag) and BOLD S1-seeded correlation map are shown on the far right. For each CAP or CDAP, the spatial similarity with regard to the correlation map is shown in the bottom right corner of each image. It can be seen that the spatial similarity increases very quickly as more frames are included, and reaches a plateau near 1 when a certain amount of frames are included. Even if only 10%∼20% of the dataset is used, most CAPs and CDAPs can replicate a spatial pattern nearly identical to the correlation map, which is calculated from the entire dataset (for LFP-CDAPs it takes a little longer, but 30% is enough to replicate the spatial pattern). It is worth noting that the display range of each image is different. Each image is normalized by its 98 percentile. The 98 percentile of LFP-BOLD correlation under ISO, BOLD S1-seeded correlation under ISO, LFP-BOLD correlation under DMED and BOLD S1-seeded correlation under DMED are 0.171, 0.299, 0.134 and 0.385 respectively. The similarities between LFP-BOLD correlation map and BOLD S1-seeded correlation map are 0.827 and 0.925 under ISO and DMED, respectively. The inter-anesthesia similarities are 0.761 and 0.569 for LFP-BOLD correlation map and BOLD S1-seeded correlation map, respectively.

**Figure A.7.**
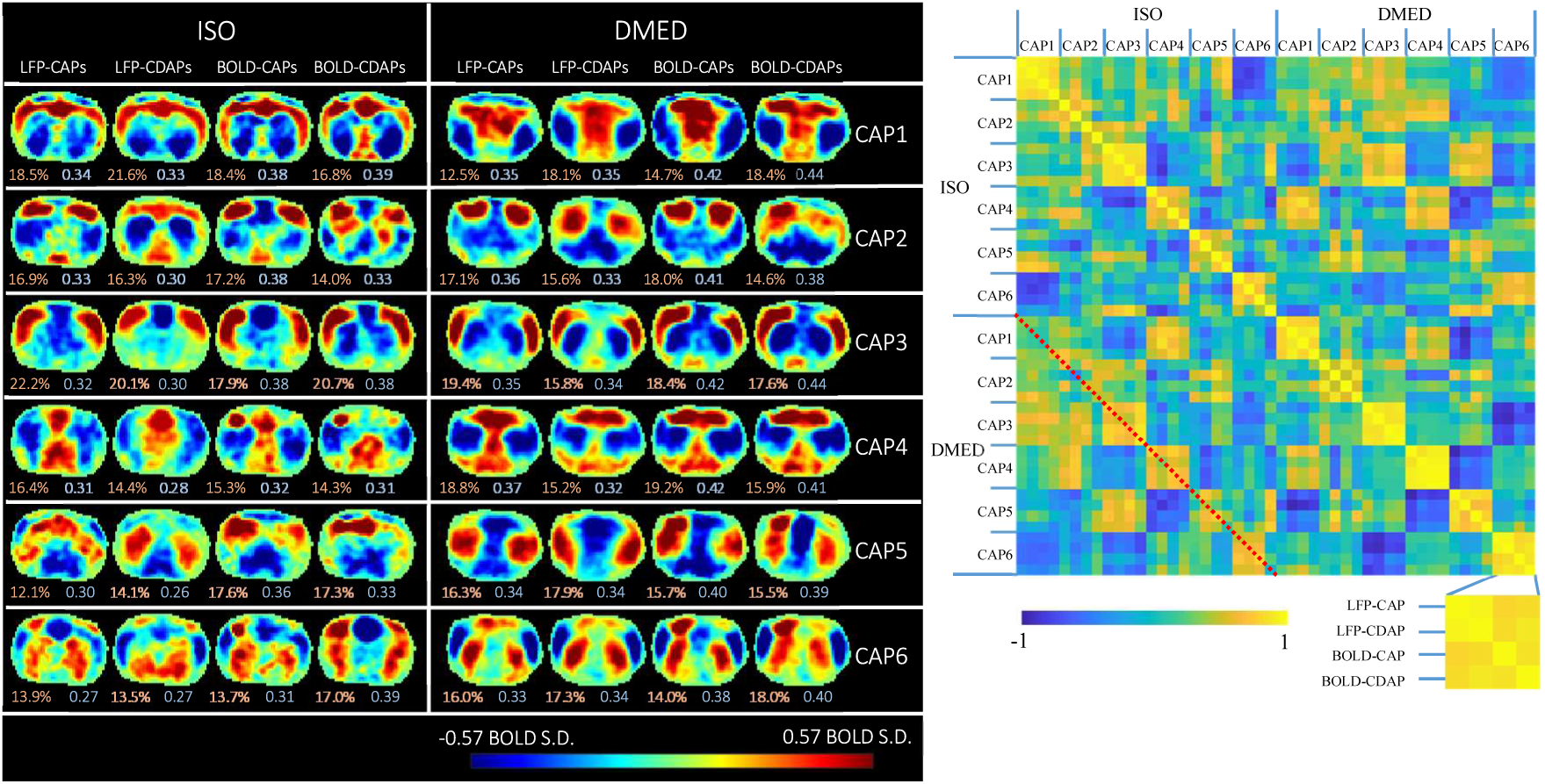
Temporal decomposition of the selected co-activation frames and co-deactivation frames. The selected frames that resemble the spatial pattern in cross correlation map are further divided into six clusters using k-means clustering (k=6) to produce CAPs and CDAPs. The threshold used for selecting frames was 15% for LFP-CAPs, BOLD-CAPs, BOLD-CDAPs, and 30% for LFP-CDAPs (the BOLD-CAPs and BOLD-CDAPs only need 5% to resemble the pattern, but to avoid randomness caused by extremely small sample size, the threshold was set to 15% as well. The LFP-CAPs under ISO are sorted based on the consistency (the average spatial similarity of each fMRI frame to the group mean). The LFP-CAPs under DMED are sorted to maximize the summed spatial similarity between LFP-CAPs under ISO and LFP-CAPs under DMED (for easier comparison across different anesthesia). The LFP-CDAPs, BOLD-CAPs, BOLD-CDAPs are also sorted in a similar way using LFP-CAPs as the benchmark. The consistency (light red) and fraction values (light blue) of the CAPs or CDAPs are shown on the bottom of each image.

